# Identification of drug candidates for rescue of SOX17 gene targets in pulmonary arterial hypertension

**DOI:** 10.64898/2026.05.14.725284

**Authors:** Eleni Vasilaki, Bedia Akosman, Shanshan Song, Rachel Walters, Yamini Sharma, Mandy Pereira, Merve Keles, Nadiya V Mykytyuk, Hannah Maude, Navneet Singh, George Field, Corey E. Ventetuolo, Luke S Howard, Jurjan Aman, Martin R Wilkins, James R. Klinger, Lan Zhao, Inês Cebola, Olin D. Liang, Christopher J Rhodes

## Abstract

**Background:** Both rare and common variants in the SRY-Box Transcription Factor 17 (*SOX17*) locus are associated with pulmonary arterial hypertension (PAH). SOX17 dysregulation leads to pulmonary artery endothelial cell (PAEC) dysfunction and the obstructive remodelling that characterises PAH.

**Hypothesis:** Impaired *SOX17* expression contributes to the pathogenesis of PAH. Restoring the function of SOX17 or its downstream targets using compounds that mimic its transcriptomic signature will rescue PAEC dysfunction and prevent PAH development.

**Methods and Results:** We defined thousands of genes with direct SOX17 genomic binding sites and identified important potential binding partners, including ETS-transcription factors such as ERG by ChIP-seq in PAECs. Through the integration of three PAEC RNA-seq datasets involving overexpression and silencing of *SOX17*, we defined a robust SOX17 transcriptomic signature. In PAH patients, circulating plasma protein levels of 10 SOX17 signature genes were associated with the *SOX17* common risk variants. This included *EFNB2* and *UNC5B*; knockdown of these genes altered the viability and apoptosis of PAECs in response to TNFα treatment. The drug-transcriptome database Connectivity Map (CMap) was used to predict novel potential therapeutic compounds to correct the SOX17 transcriptomic signature. Five compounds were selected for *in vitro* testing and were able to partially reinstate SOX17 target gene expression in PAECs. One compound, BX-912, was selected for *in vivo* testing as it corrected the levels of multiple target genes, including suppressing Runt-related transcription factor-1 (*RUNX1*). BX-912 blocked the development of pulmonary hypertension in mice lacking the *SOX17* enhancer associated with human disease.

**Conclusion:** We have demonstrated the therapeutic potential of targeting SOX17 in PAH through correction of its gene targets, identifying BX-912 as a lead compound with *in vivo* efficacy.

## Introduction

Pulmonary arterial hypertension (PAH) is a rare and complex disease characterized by obstructive remodelling of precapillary pulmonary arteries, driving increased pressure and resistance. This results in right ventricular hypertrophy, subsequent dysfunction and ultimately heart failure (1). Survival remains unacceptably poor with a recent meta-analysis estimating 88-90% survival at 1 year, 78-85% at 3-years and 61% at 5 years (2). While survival has been improved with advancements in available therapies, there is still an unmet clinical need for more effective PAH treatment approaches.

Human genetics is a powerful tool for identifying and prioritising pathogenic signalling pathways. The first established genetic risk factor for PAH was heterozygous loss-of-function rare variants in bone morphogenetic protein receptor type 2 (*BMPR2*), which are responsible for 70-80% of heritable PAH and 10-20% of idiopathic PAH cases (3). Since then, rare variants in 11 other causal genes have been associated with PAH (4–7). Amongst these causal genes is *SOX17*, which is implicated in the development of PAH through both rare (6) and common genetic risk variants; over 50% of patients with idiopathic PAH exhibit the highest-risk *SOX17* genotype (8).

SOX17 is a transcription factor that plays crucial roles in physiological vascular development and maintenance of homeostasis throughout development and life. Dysregulation of SOX17 leads to abnormal development of pulmonary (9) and retina (10) vasculature, the aortic root (11) and coronary arteries (12). Loss of SOX17 leads to endothelial cell dysfunction (13–16) and disrupts various pathways implicated in PAH development, including HGF/c-Met signalling pathway (13), E2F1 (14) and miRNAs delivered by endothelial cell (EC) exosomes (15). But there remains scant knowledge of the pathways downstream of SOX17, particularly those involved in PAEC dysfunction and the development of PAH that might provide druggable targets to restore SOX17 function.

In earlier work, we used a transcriptomic signature of *SOX17* downregulation (16) and harnessed the CMap database, which contains transcriptomic signatures of gene manipulations and treatments with small molecule compounds in various cell types (17), to predict compounds that can reverse the effects of a loss of *SOX17*. This identified Sirolimus and Aminopurvalanol-a, as candidates for partially reinstating PAEC gene expression (16).

Multi-omics studies are able to reveal molecular information that might not be apparent at a single omic level (18) and have played a crucial role in uncovering cellular mechanisms that contribute to the development of PAH (19,20). To more robustly identify SOX17 downstream targets we analysed and integrated three different levels of omics data examining gene expression (RNA-seq), patient plasma protein levels (SomaScan assay) and DNA binding (ChIP-seq), and characterised the functional importance of targets in PAEC *in vitro*. We then used our new SOX17 transcriptomic signature to predict novel therapeutic compounds for PAH that can restore the appropriate expression of SOX17 target genes and rescue PAEC dysfunction. Finally, we examined the efficacy of one compound in attenuating pulmonary hypertension (PH) in a mouse model of impaired SOX17 expression designed to mimic the common genetic variation associated with PAH in humans.

## Methods

Expanded methods and materials are available in the Supplementary Methods and Figures.

### Overexpression of *SOX17*

To examine the effect of *SOX17* overexpression on global gene expression and function of PAECs, adenoviral vectors were used to transduce a transgene of *SOX17* into human primary PAECs. Untitered seed stocks of recombinant adenovirus (DE1/E3) carrying a transgene of *SOX17* (Ad-h-*SOX17*, ADV-224019) and control adenoviral vector Ad-Null (1240, Vector Biolabs) were expanded in HEK293A cells and purified using the ViraBind™ Adenovirus Miniprep kit (VPK-099, Cell Biolabs). Adenoviral titration was performed with TCID50 (Median Tissue Culture Infectious Dose) assay and multiplicity of infection (MOI) was then calculated based on the TCID50/ml value in combination with the number or cells plated along with the virions required to achieve MOI 5 and 1.

### RNA-seq analysis

The effect of SOX17 overexpression on the transcriptome of PAECs was assessed by RNA-seq analysis. RNA was extracted using the RNeasy Plus Mini kit (Qiagen) based on manufacturer’s instructions. Sample quality was assessed and libraries were prepared based on a standard RNA-sequencing mRNA stranded protocol for the specific sequencing platform. Sample libraries were then individually barcoded, pooled and sequenced in an Illumina HiSeq 4000 with 75bp paired-end reads. Following sequencing demultiplexing was performed using the Illumina Bcl2Fastq software (21). Salmon (v1.2.1) (22) was used to align reads to GENCODE release 28 transcript annotations and calculate the abundance estimates for each transcript. Following this, the abundance estimates of the different transcripts of each gene were transformed to individual gene expression data using the R package Tximport (23). Principal components analysis was performed and PC1 and PC2 were used as covariates in all differential expression analyses to correct for variability between donors and sample quality. Differential expression analysis was performed using the Bioconductor software package edgeR (24). Adjusted p-values were calculated by edgeR using the Benjamini-Hochberg FDR method. All analyses were performed in R (version 4.2.1) (25) and RStudio (version 2022.2.3.492) (26), and volcano plots were generated using the R package ‘EnhancedVolcano’ (27). To examine the enrichment of genes participating in specific biological processes and pathways within the differentially expressed genes derived from the RNA-seq analysis an over-representation analysis was performed using the web-based Gene SeTAnaLysis Toolkit (WebGestalt).

### ChIP-seq analysis

The genome-wide binding profile of SOX17 in human primary PAECs was determined by ChIP-seq. Samples were submitted and processed by the FactorPath™ ChIP-seq service of Active Motif. Data were analysed following ENCODE best practises. Gene and genomic annotation of the SOX17 peaks were performed using the tools Genomic Regions Enrichment of Annotations Tool (GREAT) (28) and ChIPseeker (29), respectively. To examine the enrichment of transcription factor binding motifs in the SOX17 peaks a motif enrichment analysis was performed using Hypergeometric Optimization of Motif EnRichment (HOMER) (findMotifsGenome.pl) (30). To visualise the location and characterise the activity of the SOX17 binding sites on targets of interest, SOX17 ChIP-seq peaks were integrated with the GeneHancer track collection (31), which includes annotations of regulatory regions and interactions between regulatory elements, using the UCSC genome browser (32).

### Construction of a robust SOX17 signature

To facilitate the identification of novel targets of SOX17 we constructed a robust transcriptomic signature of SOX17 by combining the differentially expressed genes following SOX17 manipulation, including: a) knockdown of *SOX17* using siRNA, and b) adenoviral overexpression of *SOX17* with two MOIs (1 and 5), both validated through increased protein levels. Datasets were analysed using edgeR and the results were overlapped in R. The criteria for genes to be included in the SOX17 signature are presented in **Supplemental Table 6**.

### Downregulation of *UNC5B* and *EFNB2*

To assess the effect of the newly identified downstream targets of SOX17 we reduced their expression using silencer select siRNA (ThermoFisher) targeting *UNC5B* (s47700) and *EFNB2* (s4513) along with a negative control scrambled siRNA (negative control No.2, 4390847, Thermo Fisher Scientific).

### Cellular Function Assays

The effect of *UNC5B* and *EFNB2* downregulation, of BX-912 treatment and of *SOX17* downregulation in combination with BX-912 treatment on the function of human primary PAECs was assessed via proliferation (MTT and MTS), viability, and apoptosis (Caspase) assays as described in the **Supplementary Methods**.

### Prediction of novel therapeutic compounds using the Connectivity Map

The SOX17 signature was used to query the CMap database v1.0 (33,34) (see **Supplementary Methods**) to predict novel compounds that could reinstate SOX17 target gene expression and rescue PAEC dysfunction. Compounds were chosen based on the criteria described in the **Table 1** legend and the **Supplementary Methods**, which included consideration of our published CMap analysis that identified compounds able to correct the gene signature associated with *SOX17* promoter repression by CRISPR-mediated inhibition (16).

**Table 1.** Drug compounds predicted via CMap to reinstate SOX17 molecular signature. Five compounds were selected for further investigation (i) four based on high Tau in current search and <-70 in published SOX17 promoter repression data (ii) one, KU-0063794, based on <-90 in published SOX17 promoter repression data and high score in current search.

| **Compound ability to alter SOX17 molecular signature (Tau score)** |  | **Compound Name** | **Compound family** |
| --- | --- | --- | --- |
| MOI5 FDR<br>q<0.05, and<br>MOI1/siSOX17<br>p<0.05, | SOX17<br>Promoter<br>Repression by<br>CRISPR,<br>Walters et al. |  |  |
| **97.57** | **-90.95** | linifanib | PDGFR inhibitor |
| **93.8** | **-81.25** | PIK-90 | PI3K inhibitor |
| **93.78** | **-70.18** | TG-101348 | FLT3 inhibitor |
| **93.35** | **-84.22** | BX-912 | 3-Phosphoinositide-dependent protein kinase-1 (PDK1) inhibitor |
| 87.86 | **-91.56** | KU-0063794 | MTOR inhibitor |
| 90.19 | -72.9 | PP-30 | RAF inhibitor |
| 88.74 | -54.34 | PI-828 | PI3K inhibitor |
| 90.52 | -17.82 | GW-583340 | EGFR inhibitor |
| 98.13 | 0 | FR-180204 | Not specified |
| 96.26 | 0 | PF-562271 | Focal adhesion kinase inhibitor |
| 96.01 | 0 | colforsin | Adenylyl cyclase activator |
| 95.74 | 0 | chromomycin-a3 | DNA binding agent |
| 93.82 | 0 | 1-phenylbiguanide | Serotonin receptor agonist |
| 91.09 | 0 | ZG-10 | JNK inhibitor |

The effect of the candidate compounds on gene expression in human primary PAECs was evaluated at different concentrations (10µM, 3 µM, 1µM and 100nM as per CMap) after 6 and 24 hours. For each compound, the expression levels of four to six genes consistently predicted to change by CMap was measured.

We then calculated the ‘net benefit percentage’, which captures the proportion of genes regulated in the predicted direction and subtracts those genes regulated in the opposite direction for each compound across all treatment concentrations and timepoints.

### Reverse Transcription - Quantitative PCR

Changes in expression of target genes were measured with reverse transcription–quantitative polymerase chain reaction (RT-qPCR) with β-actin (*ACTB*) as the reference gene.

### Protein quantification by Western Blot

Cells were lysed with a mixture of RIPA buffer (Sigma) with Halt™ Protease Inhibitor Cocktail (Thermo Fisher Scientific) and phosphatase inhibitors (Sigma) following incubation on ice for 3 minutes and then centrifugation at 4°C for 5 minutes at max speed to pellet any cellular debris.

Protein quantification of samples was performed with the Pierce™ BCA Protein Assay (Thermo Fisher Scientific) and protein concentrations were calculated using a standard curve.

Proteins were diluted in Laemmli SDS sample buffer (4X, Thermo Fisher Scientific) and denatured at 95°C for 5min prior to being loaded in NuPAGE™ 4 to 12%, Bis-Tris gels (Thermo Fisher Scientific) along with the Spectra™ Multicolor Broad Range Protein Ladder (Thermo Fisher Scientific) and run at 170V for 45min in in NuPAGE MES-SDS running buffer (1X, ThermoFisher). Transfer of the protein bands was performed using the Bio-Rad Trans-Blot Turbo Transfer System, after which PVDF membranes were incubated in 5% milk in TBST for 1 hour at RT to block non-specific binding. Primary antibodies were diluted at the given concentrations in 5% BSA in TBST (**Supplemental Table 5**) and incubated overnight at 4°C, while secondary antibodies were diluted in 5% milk in TBST and incubated at RT for 1 hour. Membranes were washed between antibody incubations with TBST. Bands were visualised using the Immobilon Crescendo Western HRP substrate and a ChemiDoc (Bio-Rad) device paired with ImageJ (35) software (version 1.53e).

### Animal models of PH in *SOX17* enhancer knock out and BX-912 treatment

Transgenic mice lacking the *SOX17* signal 1 region (*SOX17* enhancer knockout, SOX17enhKO) develop severe PH in response to mild Sugen hypoxia (mSuHx, 12% O2, 5 mg/kg Sugen-5416) as described in (16). In the disease intervention protocol, treatment with BX-912 was started one week after the mSuHx; mice were given 50 mg/kg of BX-912 in 100 µl of DMSO, or 100 µl vehicle alone by oral gavage every other day, for a total of six doses. Right ventricular systolic pressure (RVSP) and right ventricular (RV) hypertrophy (RV to left ventricle + septum wet weight (RV/LV+S) ratio) were measured along with immunohistochemical staining (IHC) of mouse α-smooth muscle actin to assess vessel muscularisation. Details of these procedures are provided in the Supplementary Methods.

### Statistical analysis

Statistical analysis and data visualization was performed using GraphPad Prism 10 (GraphPad Software, San Diego, California, USA), R (version 4.2.1) (25) and RStudio (version 2022.2.3.492) (26). Data are presented as mean ± standard deviation (SD). Normally distributed data were analysed by parametric methods, with transformations applied to normalize where required, and specific tests listed in the figure legends. Plasma proteomics data analyses were detailed previously (16).

Relative gene expression values were normalised based on the average of all 2^ΔCt values within each experiment (**Figure 2B**) or within each independent condition (**Figure 4B**, **Figure 5A**) to account for variability between experiments.

**Table 2.**
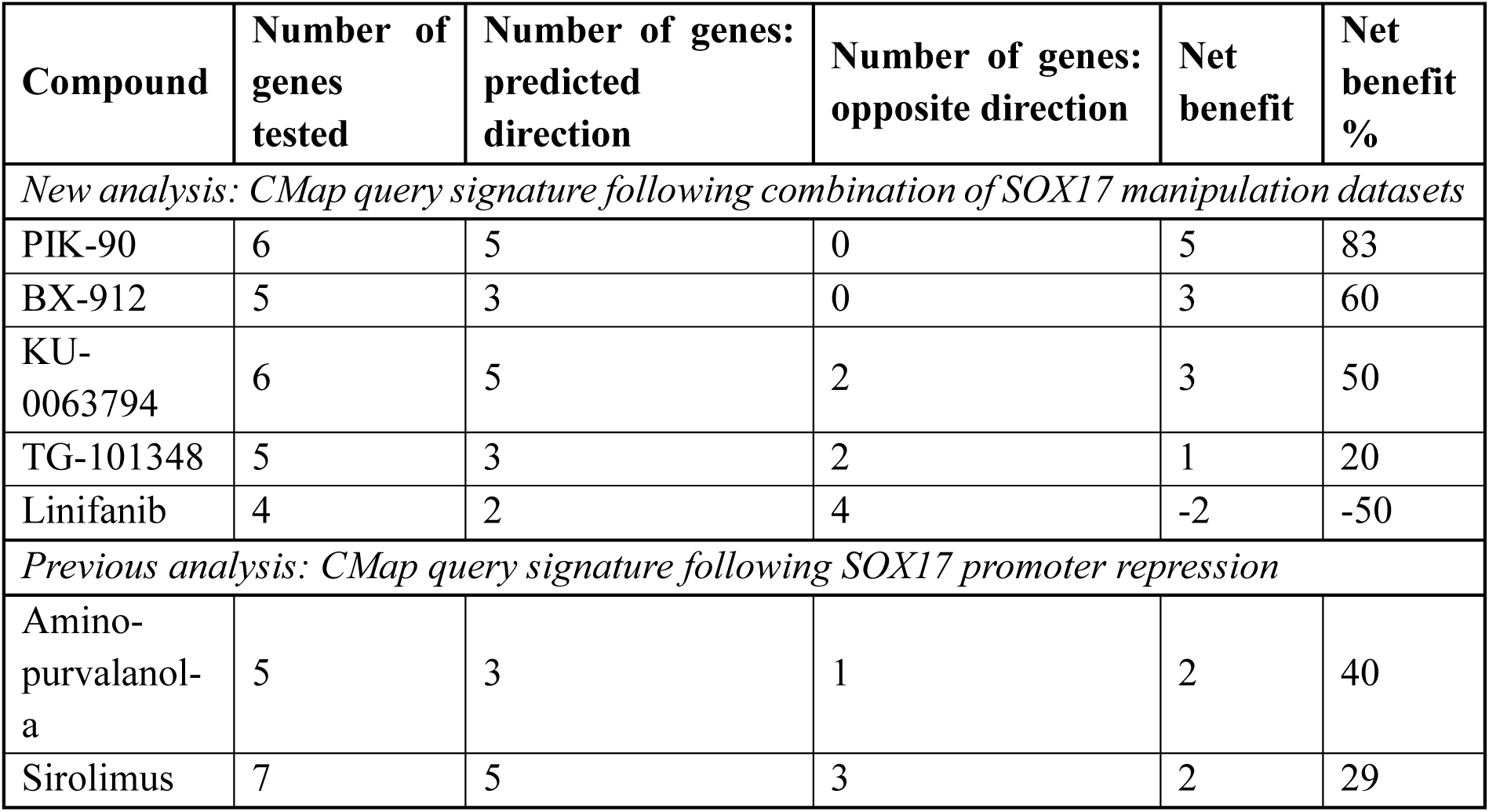
Effectiveness of the novel CMap-predicted SOX17-activity restoring compounds. Net benefit was calculated based on the number of genes tested and the number of the conditions tested where gene expression was significantly changed in the predicted or opposite direction. This was also calculated for the compounds predicted in our previous publication for reference.

| **Compound** | **Number of genes tested** | **Number of genes: predicted direction** | **Number of genes: opposite direction** | **Net benefit** | **Net benefit %** |
| --- | --- | --- | --- | --- | --- |
| *New analysis: CMap query signature following combination of SOX17 manipulation datasets* |  |  |  |  |  |
| PIK-90 | 6 | 5 | 0 | 5 | 83 |
| BX-912 | 5 | 3 | 0 | 3 | 60 |
| KU-0063794 | 6 | 5 | 2 | 3 | 50 |
| TG-101348 | 5 | 3 | 2 | 1 | 20 |
| Linifanib | 4 | 2 | 4 | -2 | -50 |
| *Previous analysis: CMap query signature following SOX17 promoter repression* |  |  |  |  |  |
| Amino-purvalanol-a | 5 | 3 | 1 | 2 | 40 |
| Sirolimus | 7 | 5 | 3 | 2 | 29 |

For the protein quantification (**Figure 2C**, **Figure 5B**) the intensity of the protein bands was measured using the ImageJ software and then normalised to β-ACTIN.

PAEC functional assay (**Figure 3**, **Figure 5C**) absorbance and luminescence values were corrected for blank wells and normalised based on the average of all corrected values within each experiment to account for variability between experiments. The apoptosis/viability ratio was calculated using the corrected and normalised values for apoptosis and viability for each sample.

PAEC MTS assay absorbance values (**Figure 5D&E**) were corrected for the background absorbance (medium and MTS reagent without cells).

## Results

### SOX17 directly regulates a plethora of genes involved in crucial EC functions

#### ChIP-seq identification of SOX17 genomic binding targets in PAECs

In order to identify direct downstream targets of SOX17, we examined the genome-wide binding of SOX17 in human primary PAECs by ChIP-seq. 15,302 replicated peaks were defined across four biological samples, and they were evenly distributed between promoters (22.02%) and genomic regions where enhancers are usually found (intronic regions apart from the 1st intron (29.63%), and distal intergenic regions (25.53%) **Figure 1A**).

**Figure 1.**
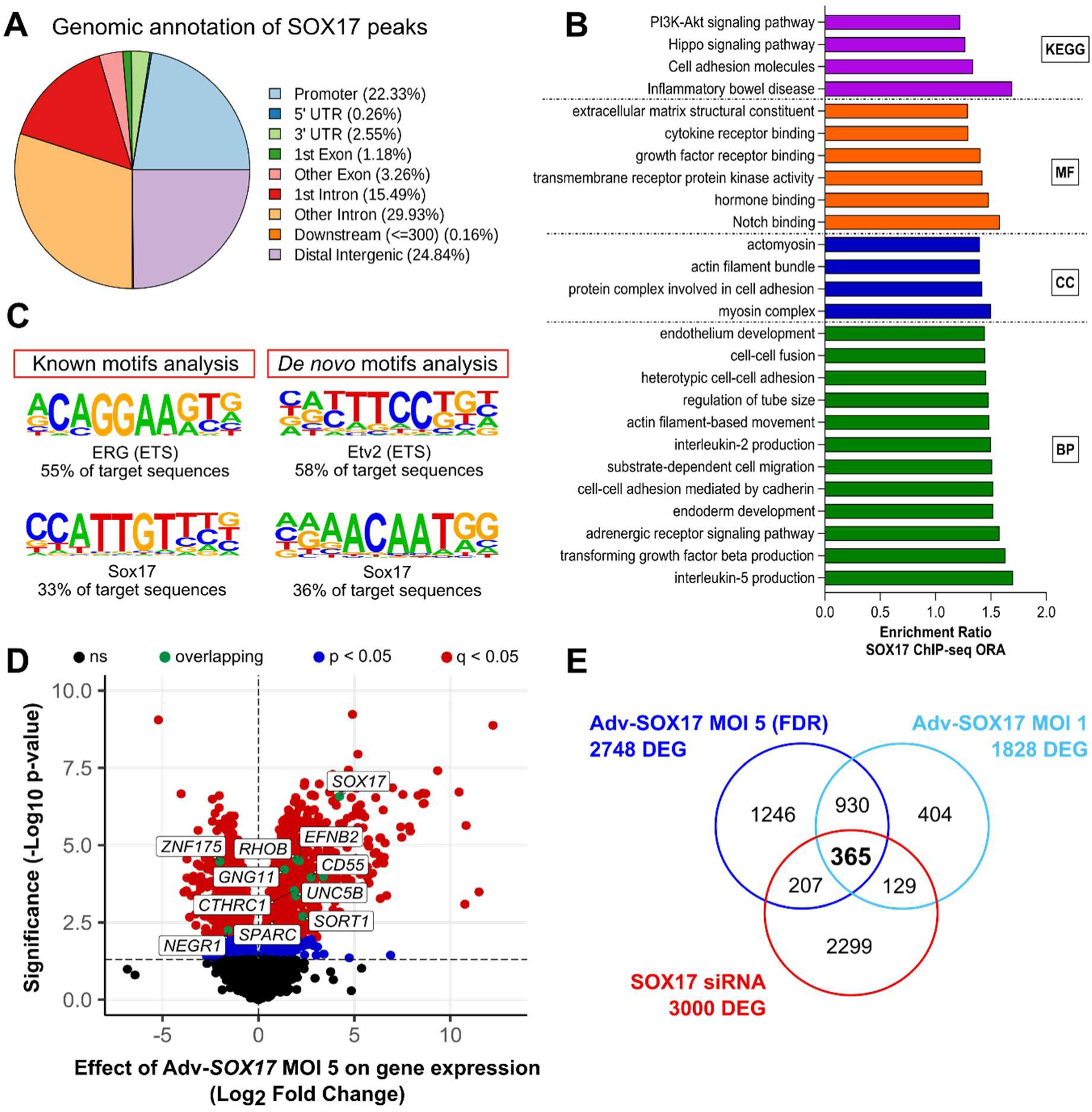
SOX17 directly binds and regulates genes with PAEC relevant functions. **A)** Distribution of SOX17 peaks based on their genomic annotations. **B)** ORA of the genes bound by SOX17; biological process (BP), cellular component (CC), molecular function (MF), Kyoto Encyclopedia of Genes and Genomes pathway (KEGG), q < 0.05 for all categories presented. **C)** Motif enrichment analysis on the SOX17 peaks to uncover potential co-factor. **D)** Differential gene expression in PAECs following SOX17 o/e (MOI 5, 48 hours). The labelled green overlapping genes refer to the genes commonly identified in all three omics shown in Figure 2A) Overlap between RNA-seq datasets following SOX17 manipulation (siRNA-SOX17, SOX17 o/e MOI 5 (FDR) & MOI 1) to uncover genes directly regulated by SOX17. For details on the thresholds used to filter the datasets see **Supplemental Methods** and **Supplemental Table 6**.

To annotate putative target genes, we used GREAT, which identified 6,236 genes after filtering for expression in PAECs (**Supp Excel 2 - GREAT gene annotation.xlsx**). These putative targets are enriched for genes associated with biological processes, molecular functions, and pathways relevant to endothelial function and PAH, including TGF-β production, endothelial development, extracellular matrix organisation, Notch signalling, and cell adhesion (**Figure 1B**).

The function of SOX family members is significantly affected by co-factors. Therefore, we examined whether the PAEC SOX17 binding sites (200 bp in the centre of peaks) are enriched for binding motifs of other transcription factors (TF), as those may act as co-factors of SOX17. The most enriched known motifs detected correspond to ERG (55.05% of target sequences, 17.34% of background, p=10^-2409^, **Figure 1C**). ERG is a member of the Erythroblast Transformation Specific (ETS) family and four other ETS family TFs were enriched in ranks 2 to 5, reflecting the high degree of similarity of binding motifs across this TF family.

Besides the ETS binding motifs (9 out of the top 20 matches), there was also enrichment of motifs belonging to Basic Leucine Zipper Domain (bZIP) TFs such as FOS and FRA1, accounting for 8 out of the top 20 matches (**Supp Excel 4 - Primary & coenriched motifs.xlsx**). The *de novo* motif analysis indicated enrichment of binding sites of the TF ETV2, another member of the ETS family (58.03% target, 18.69% background p=10^-2525^), SOX17 (35.58% target, 8.3% background, p=10^-1930^), bZIP JunB (25.49% target, 3.95% background, p=10^-1904^) and GATA4 (19.61% target, 7.6% background, p=10^-494^, **Figure 1C and Supp Excel 4 - Primary & coenriched motifs.xlsx**). These results suggest that SOX17 may cooperate with ETS and bZIP TFs for the regulation of endothelial cell gene expression. In addition, the analysis identified the binding of TFs cooperating with β-catenin, a known co-activator of SOX17, such as TCF4 (19.15% target, 13.78% background, p=10^-75^) and LEF1 (8.7% target, 7.01% background, p=10^-14^).

#### Transcriptomic analysis of SOX17 targets in PAECs

To further characterise SOX17 downstream targets, we examined the effect of manipulating SOX17 levels on the global gene expression of PAECs. We have previously shown that knockdown of *SOX17* via siRNA in PAECs resulted in the differential expression of 1717 genes (16). We now examined the effects of overexpressing *SOX17* in PAECs using adenoviral vectors at two different viral loads (MOI 5 and MOI 1) on gene expression. These produced a 19-fold and 7-fold increase in *SOX17* mRNA at 48 hours (**Supplementary Figure 1**), and both produced a significant increase in protein levels (**Figure *2*C, Supplementary Figure 5**). The higher MOI resulted in the differential expression of 2748 genes (MOI 5, q<0.05 & Log_2_FC > |0.25|, **Figure *2*D**, **Supp Excel 1 - SOX17 RNAseqs DEG.xlsx**). 1295 out of the 2748 significantly differentially expressed genes (DEG) with MOI 5 were also nominally significant and directionally consistent with the lower MOI (MOI 1, p<0.05 & Log_2_FC>|0.25|, **Supp Excel 1 - SOX17 RNAseqs DEG.xlsx and Supplementary Figure 2**). Differentially expressed genes following *SOX17* overexpression were enriched for processes and pathways implicated in the development of PAH, such as vascular development and angiogenesis, regulation of cell adhesion, cell junction organisation and extracellular structure organization (**Supplementary Figures 3&4**).

**Figure 2.**
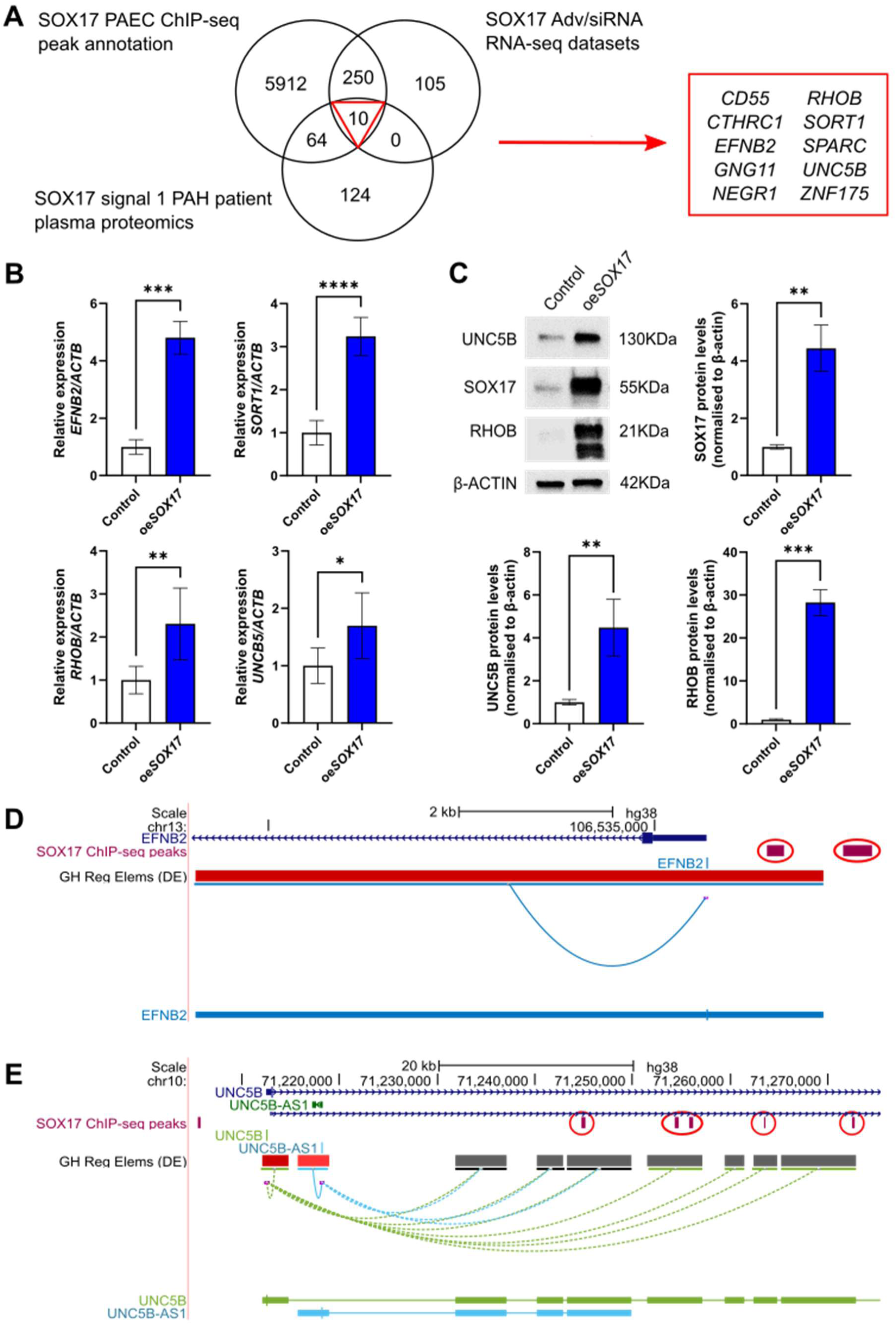
Integration of multi-omics data leads to the identification of novel direct downstream targets of SOX17 in PAEC. **A)** Overlap of genes identified by SOX17 ChIP-seq gene annotation, RNA-seq following SOX17 manipulation and proteomics analysis in the plasma of PAH patients based on SOX17 enhancer signal 1 genotype. **B)** Relative expression of novel downstream targets following SOX17 o/e with adenoviral vectors at MOI 5 (from now on Adv-*SOX17* MOI 5 referred as oe*SOX17* and Adv-Null MOI 5 referred as Control) for 48 hours in PAECs (*EFNB2* n=4, *SORT1* and *UNC5B* n=7, Welch’s t test, *RHOB* n=7, Mann-Whitney test, relative values calculated based on average of Adv-C (MOI 5 and 1 separately) from all experiments). **C)** Protein levels of SOX17 and downstream targets following SOX17 o/e with MOI 5 for 48 hours (oe*SOX17*) in PAECs (relative values calculated based on the average of Adv-C (MOI 5 and 1 separately) from all experiments and normalised by log10-transformation, n=3, Welch’s t-test, relative values presented). Binding of SOX17, as indicated by the replicated peaks of the SOX17 ChIP-seq analysis in PAECs, near the TSS and within putative enhancer regions of **D)** *EFNB2*, and **E)** *UNC5B*. Binding regions of SOX17 presented as purple bars and circled in red. Regulatory regions annotation and TSS provided by the GeneHancer collection; double elite (DE) means that elements are derived by more than one database included in the collection. TSS are indicated as lines underneath the SOX17 ChIP-seq track. Regulatory element annotation is presented underneath the TSS; dark red= high confidence promoter, medium red= medium confidence promoters, dark grey= high confidence enhancers. Next track indicates the interactions between regulatory elements; dashed line indicates that the TSS precedes the interacting element. Last track presents the clustered interactions between regulatory elements of each gene. *p<0.05, **p<0.01, ***p<0.001, ****p<0.0001

To prioritise robust targets of SOX17, we integrated the results from our previous *SOX17* knockdown experiments (16) with the two adenoviral-mediated overexpression results, identifying 365 DEG (‘SOX17-DEG’) (**Figure 2E and Supp Excel 1 - SOX17 RNAseqs DEG.xlsx**) in common and demonstrating a significant overlap (p<0.0001, Fisher’s exact test). This list of SOX17-DEG was taken forward to query the CMap database to predict novel potential therapeutic compounds for PAH. Moreover, we aimed to further validate and characterise a subset of these DEG with patient data and *in vitro* functional studies.

### Regulation of novel SOX17 targets is validated in PAH plasma proteome and *in vitro*

#### SOX17 targets in patient plasma proteomic profiles are determined by SOX17 genotype

We previously showed that disease risk variants at *SOX17* enhancer signal 1 significantly affect the levels of 198 proteins in the plasma of PAH patients (16). Of the 260 genes that are both regulated and bound by *SOX17* (**Figure 2A**), 73 were measured in the proteomic analysis. Of these, 10 were associated with the *SOX17* signal 1 enhancer genotype, including five with known EC functions: Unc-5 Netrin Receptor B *(UNC5B),* Collagen Triple Helix Repeat Containing 1 *(CTHRC1),* Ephrin B2 *(EFNB2),* Ras Homolog Family Member B *(RHOB)* and Sortilin 1 *(SORT1,* **Figure 2A**).

#### Validation of the regulation of novel downstream targets by SOX17

To validate the effect of SOX17 on the gene expression and protein levels of these five genes in PAECs, we performed RT-qPCR and Western blot analyses after adenoviral overexpression, observing significant increases in the expression of *EFNB2, SORT1, RHOB* and *UNC5B* expression (**Figure 2B, Supplementary Figure 5A**), consistent with the RNA-seq data (**Supp Excel 1 - SOX17 RNAseqs DEG.xlsx**). There was no apparent effect on the expression of *CTHRC1* (data not shown).

Immunoblot analysis further confirmed that both MOIs of the adenoviral vector significantly increased in SOX17 protein levels and that SOX17 overexpression led to significant increases in protein levels of RHOB and UNC5B (**Figure 2C, Supplementary Figure 5B**).

#### Examination of SOX17 binding at novel target genes

We also examined the functionality of the regions bound by SOX17 near the candidate target genes by combining the SOX17 ChIP-seq peaks with public datasets containing annotations of regulatory regions and interactions between regulatory elements (see **Supplementary Methods**). Based on the GeneHancer Regulatory Elements track, which annotates promoters in red and enhancers in grey with darker colours indicating higher confidence score (36) SOX17 binding was present within the promoter of *EFNB2*, both upstream and near the transcription start site (TSS) of the gene (**Figure 2D**). A second peak was present slightly upstream of the first and outside of the promoter region (**Figure 2D**). *SORT1* presented similarly, with binding of SOX17 near the TSS and within the promoter region (**Supplementary Figure 6A**). For *RHOB* there was binding of SOX17 at the TSS but also multiple binding sites in a distal intragenic region, which was annotated as a strong enhancer interacting with the TSS of *RHOB,* based on genomic annotations and element interaction marks (**Supplementary Figure 6B**).

The landscape of *UNC5B* presented a more complex pattern; one binding site of SOX17 was upstream of the promoter, but the region has no annotation based on public epigenomic datasets. However, there were five intronic binding sites of SOX17 in *UNC5B*. These regions are annotated as strong enhancers and likely interact with the promoter of *UNC5B* (**Figure 2D**). *CTHRC1* also had *SOX17* binding sites in both the promoter and enhancer regions (**Supplementary Figure 6C**).

### SOX17 and its downstream targets control PAEC function

#### Effect of SOX17 overexpression on PAEC function

We next examined the effect of SOX17 manipulation on PAEC function. SOX17 overexpression significantly reduced PAEC proliferation in response to VEGF (p=0.0012, **Figure 3A)**, as well as significantly increased apoptosis in response to TNF-α. The latter was not simply driven by altered cell density, as cell viability was decreased under the same conditions, resulting in a significantly increased apoptosis/viability ratio (all p<0.001 and **Figure 3B-D**, respectively).

**Figure 3.**
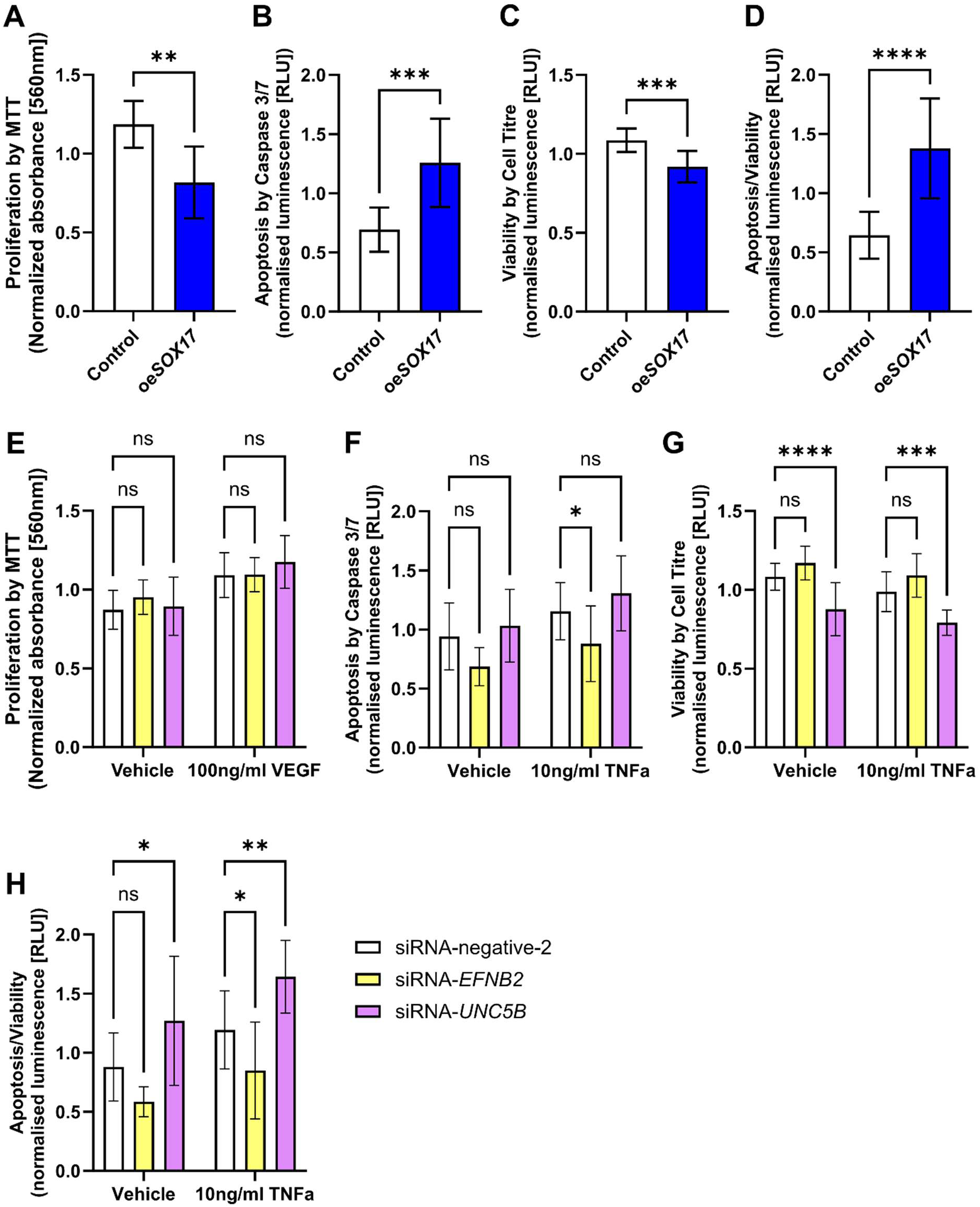
Effect of *SOX17* overexpression and targets *EFNB2* and *UNC5B* knockdown on PAEC proliferation, apoptosis and viability. Functional assays following *SOX17* overexpression with adenoviral vectors at MOI 5 (oe*SOX17*) or downregulation of *EFNB2* and *UNC5B* via siRNA for 48 hours. **A)** PAEC proliferation was assessed via MTT (3 independent experiments, unpaired t-test with Welch’s correction) after 24-hour treatment with VEGF (100 ng/ml), **B-D)** PAEC apoptosis assessed via Caspase 3/7 assay and viability assessed via Cell Titre Glo assay (4 independent experiments, unpaired t-test with Welch’s correction) after 24 hour treatment with TNF-a (10 ng/ml, for apoptosis and viability). **E)** PAEC proliferation was assessed via MTT after 24-hour treatment with VEGF (100 ng/ml), **F-H)** PAEC apoptosis assessed via Caspase 3/7 assay and viability assessed via Cell Titre Glo assay after 24-hour treatment with TNF-a (10 ng/ml, for apoptosis and viability) (5 independent experiments, 2-way ANOVA, Šídák’s multiple comparisons test). *p<0.05, **p<0.01, ***p<0.001, ****p<0.0001

#### Novel downstream targets of SOX17 affect PAEC function

To determine whether the observed effects of SOX17 on PAEC function could be mediated through any of the newly identified targets, we examined the effect of *UNC5B* or *EFNB2* knockdown (**Supplementary Figure 7**) on PAEC proliferation, apoptosis and viability. Loss of either gene did not affect PAEC proliferation (**Figure 3E**) but significantly affected apoptosis and viability (**Figure 3F-H**). Downregulation of *UNC5B* led to a significant decrease in PAEC viability (p≤0.0001, **Figure 3G**), independently of the treatment, with an increase in apoptosis normalised to viability (p=0.0146 in vehicle treatment and p=0.0035 in TNFα treatment) (**Figure 3H**). In contrast to our hypothesis, downregulation of *EFNB2* led to decreased apoptosis that was statistically significant under TNFα treatment (p=0.0326, **Figure 3F**), without reducing PAEC viability (**Figure 3G**). This resulted in a significant reduction in apoptosis normalised to viability (p=0.0404 in TNFα treatment, **Figure 3H**), the opposite effect to that observed following loss of SOX17 (16). This suggests that the downstream effects of SOX17 on PAEC function could be mediated by regulation and function of UNC5B in balance with contrasting effects of other targets including EFNB2.

### Novel therapeutic compounds for PAH predicted through robust SOX17 transcriptomic signature

Given the observations described above that perturbing SOX17 levels leads to changes in the expression of genes involved in cellular and molecular processes relevant for PAH pathogenesis, we hypothesised that restoring the expression of SOX17 downstream targets could present a novel therapeutic avenue for PAH. To approach this, we queried the compound-gene signature database, CMap, to predict compounds that can correct expression of our SOX17-DEG and thereby potentially rescue PAEC dysfunction (**Figure 4A**). From the CMap-predicted compounds, four had a strong positive correlation to SOX17 activity (Tau>90) and also showed a strong negative correlation with *SOX17* repression identified in our previous CMap query, which was based on CRISPRi targeting the *SOX17* promoter (16). Moreover, one compound had a Tau∼88 and also a stronger negative correlation (Tau -92) in the previous query (16) (**Table 1**). These five compounds were taken forward for *in vitro* validation of gene expression changes in PAECs.

**Figure 4.**
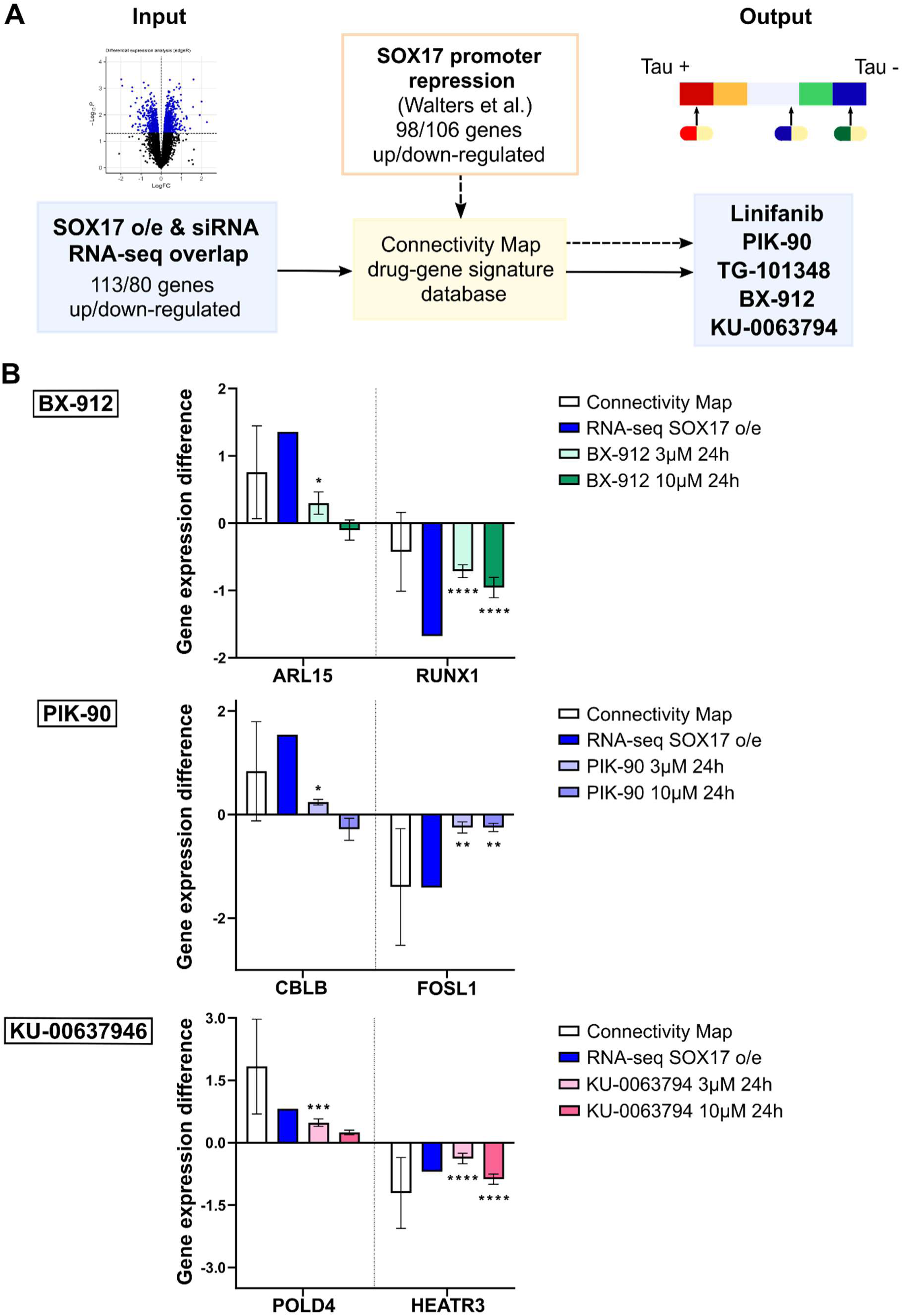
Using the robust SOX17 transcriptomic signature to predict novel therapeutic compounds. **A)** Flow-chart illustrating use of CMap database to analyse our SOX17 target gene signatures to rank drugs by Tau score (ranging from 100 to -100), which estimates how similar or opposite a drug signature is to the input signature. Note that: i) a Log_2_FC > |0.35| was used to further reduce the query gene list due to CMap input limits (see **Supplemental Methods** and **Supplemental Table 2** for details); ii) not all genes are measured in CMap so gene numbers here reflect final used gene lists. **B)** Effect of top three most effective compounds (PIK-90, BX-912, KU-0063794) treatment on selected SOX17 target gene expression in the CMap and independent experiments in PAEC, with SOX17 effects shown for reference. Average Z-scores derived from CMap database for drug target genes. Fold change derived from the DEG analysis of the RNA-seq dataset following SOX17 overexpression (MOI 5). Relative gene expression in PAECs following treatment with 3µM and 10µM of indicated drug for 24 hours. Statistics show the comparison between treatment and vehicle (DMSO). 2-way ANOVA with Sidak’s multiple comparisons test. N=3, * - p< 0.05, ** - p<0.01, *** - p<0.001, **** - p<0.0001. Values represent the fold change based on the average of the DMSO control of all 3 independent experiments.

To study the effect of the chosen compounds on gene expression, we exposed human PAECs *in vitro* to the compounds and assessed predicted target genes by RT-qPCR. All compounds except TG-101348 showed no apparent gross effects on cultures, whereas TG-101348 induced cell death and thus was excluded from further experiments.

We defined response by the proportion of genes affected in the predicted versus opposite direction (‘net benefit’). The most effective compound was PIK-90 (83% net benefit in gene expression), followed by BX-912 (60% net benefit) (**Table 2**, **Figure 4B**). KU-0063794 had a net benefit of 50% with unexpected regulation observed for two genes only at the highest concentration after 24 hours. TG-101348 had a net benefit of 20%, while Linifanib showed a negative net benefit, which was mostly driven by unpredicted regulation of genes after treatment with the highest concentration for 6 hours.

With the use of the net benefit score, it also becomes apparent that the construction of a more robust profile of SOX17 targets leads to the prediction of compounds with higher effectiveness (net benefits of 83-50%). In contrast, compounds predicted in our previous study using a single transcriptomic signature following SOX17 downregulation showed lower effectiveness (net benefits at 40% and 29%) (**Table *2***) (16).

BX-912, the compound with the second-highest net benefit of 60%, significantly downregulates RUNX1 (identified as a SOX17 target in this study), an effect that has been previously shown to ameliorate PH development *in vivo* (37). In addition, BX-912 is a PDK1 (3-phosphoinositide-dependent protein kinase-1) inhibitor, and thereby prevents activation of AKT (38), a key mechanism driving vascular remodelling in PH. We therefore selected BX-912 for further evaluation as a potential therapeutic for PAH.

### BX-912 affects both SOX17 expression and PAEC function

To test whether BX-912 could rescue SOX17 function through direct regulation, we measured *SOX17* mRNA and protein levels following drug exposure in PAEC. We observed a significant increase in *SOX17* mRNA expression at 24 hours of 10µM BX-912 treatment compared to vehicle (p=0.0069) (**Figure 5A**) but no significant effect on protein levels (**Figure 5B**). Both mRNA and protein levels (p=0.0017) of the predicted BX-912 target *RUNX1* were significantly reduced following BX-912 exposure at 10µM for 24 hours (**Figure *4*B** & **Figure *5*B**, respectively).

**Figure 5.**
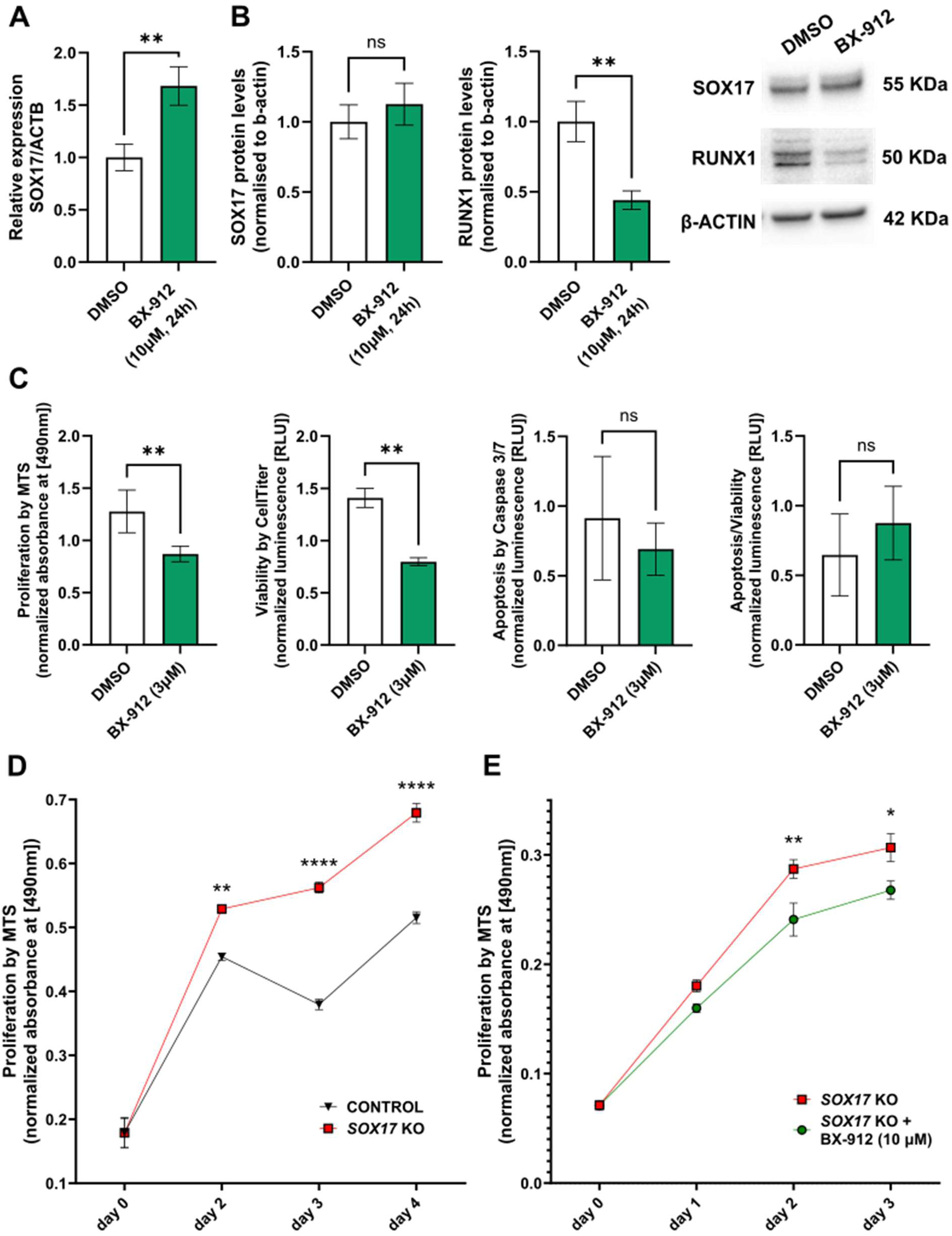
Effect of BX-912 treatment on Sox17 expression and PAEC function. **A)** SOX17 expression in PAECs following treatment with vehicle (DMSO) or BX-912 (10µM) for 24 hours by RT-qPCR (n=3, log_10_ normalised, Unpaired t test with Welch’s correction, relative values presented); **B)** Protein levels of SOX17 and RUNX1 following treatment with vehicle (DMSO) or BX-912 (10µM) at 24 hours (n=4, Unpaired t test with Welch’s correction). **C)** PAEC proliferation assessed by MTS following treatment with BX-912 (3µM) for 24 hours (n=5, Unpaired t test with Welch’s correction); PAEC viability and apoptosis assessed by Cell Titre Glow and Caspase 3/7, respectively, following treatment with BX-912 (3µM) for 24 hours (n=3, Unpaired t test with Welch’s correction); **D)** effect of SOX17 KO via CRISPR-Cas9 on PAEC proliferation assessed via MTS (ctrl n=6, SOX17 KO n=4, 2-way ANOVA with Sidak’s multiple comparisons test; **E)** effect of BX-912 treatment (10µM) on PAEC proliferation following SOX17 KO, assessed via MTS (day 0 n=4 for all conditions, rest of the days: SOX17 KO n=11, SOX17 KO + BX912 n=6, 2-way ANOVA with Sidak’s multiple comparisons test. * - p< 0.05, ** - p<0.01, *** - p<0.001, **** - p<0.0001.

We then examined whether BX-912 could alter PAEC function and rescue the effects elicited by loss of *SOX17*. BX-912 treatment for 24 hours at 3 µM led to significantly decreased PAEC proliferation (p=0.0083) and viability (p=0.0028) without affecting apoptosis or the apoptosis to viability ratio (**Figure 5C**). Moreover, BX-912 treatment inhibited the increase in proliferation caused by loss of *SOX17* via CRISPR-knockout (**Figure 5D&E**).

### BX-912 alleviates PH development in SOX17 enhancer knockout mice

In previous studies, we demonstrated that transgenic mice with deletion of *SOX17* signal 1 (SOX17enhKO), designed to mimic the effect of common risk variants associated with human PAH, develop more severe PH when exposed to hypoxia (16). These mice are also more susceptible to PH induced by Sugen hypoxia and develop severe PH when exposed to mild hypoxia and low dose Sugen-5416 (12% O_2_, and 5 mg/kg, mSuHx) which causes no PH in wild-type mice.

To determine if BX-912 is sufficient to prevent PH in SOX17enhKO mice, we administered BX-912 (50mg/kg) by oral gavage every other day, for a total of 6 doses, starting 1 week after exposure to mSuHx (**Figure 6A**). SOX17enhKO mice treated with vehicle alone developed severe PH as demonstrated by more than a doubling of RVSP and RV/LV+S (**Figure 6B&C**, respectively). Treatment with BX-912 prevented the increase in RVSP and significantly reduced RV/LV+S (**Figure 6B&C**, respectively). Concordant with this, pulmonary vascular remodelling, as evidenced by increased muscularisation of arterioles in SOX17enhKO mice exposed to mSuHx, was absent in animals treated with BX-912 (**Figure 6D and Supplementary Figure 8**).

**Figure 6.**
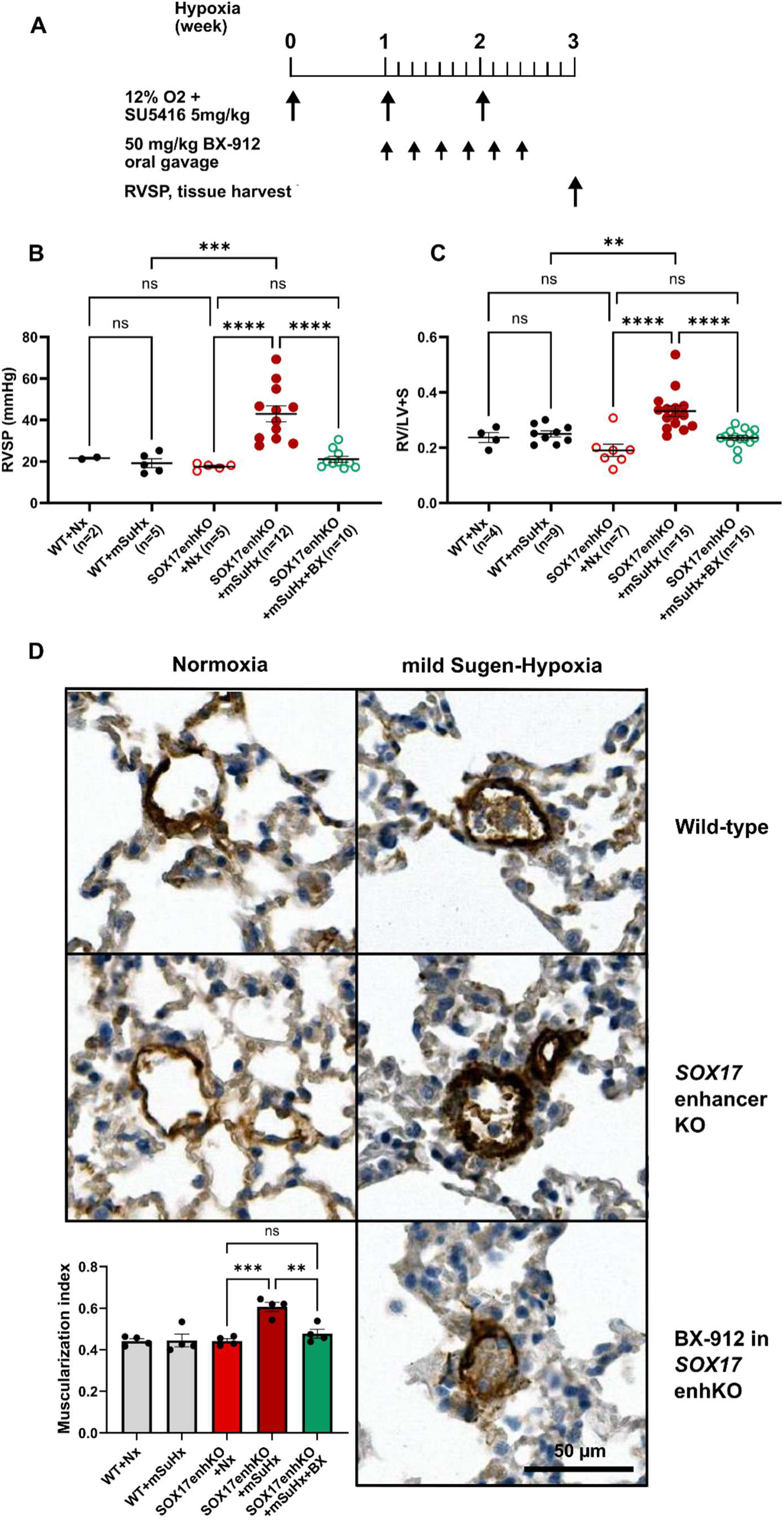
BX-912 alleviates SuHx-PH in SOX17enhKO mice. **A)** Schematic of the experiments. **B)** The mild Sugen/hypoxia (SuHx) condition (12% O_2_ and 5 mg/kg SU5416) induced pulmonary hypertension (PH) in SOX17 enhancer knockout (SOX17enhKO) mice but not in wild-type (WT) mice. The latter (WT SuHx) group had normal RVSP, which is comparable to that of the WT mice in normoxia (Nx). Without intervention, SuHx treated SOX17enhKO mice developed significantly higher RVSP than Nx treated SOX17enhKO mice. BX-912 treatment completely blocked the increase of RVSP in SOX17enhKO mice despite the SuHx treatment. **C)** Similarly, without intervention, SuHx treated SOX17enhKO mice developed significantly more severe RV hypertrophy, as indicated by increased RV/LV+S ratio. BX-912 treatment of SOX17enhKO mice completely blocked the increase of the RV/LV+S ratio despite the SuHx treatment. **D)** IHC of mouse α-smooth muscle actin for quantification of muscularisation of pulmonary arterioles <50µM. N=4 mice were included in each group, and 66 vessels were analysed in WT+Nx group, 52 in WT+mSuHx group, 60 in KO+Nx group, 115 in KO+mSuHx group and 115 in KO+mSuHx+BX group; ordinary one-way ANOVA, Tukey’s multiple comparisons test, * - p< 0.05, ** - p<0.01, *** - p<0.001, **** - p<0.0001.

## Discussion

In this study, we set out to identify new therapeutics for PAH based on the concept that correction of gene expression changes triggered by dysfunction of SOX17 in the pulmonary endothelium can improve endothelial function and thereby reverse the disease progression. Using integrated multi-omics analyses, we defined a robust SOX17 transcriptional signature, identified novel downstream targets involved in the regulation of PAEC function and differentially expressed according to *SOX17* genotype. We also identified novel therapeutic compounds that reinstate expression or repression of SOX17-associated genes in PAECs. Finally, we demonstrated that our lead compound, BX-912, inhibits PAH development in genetically predisposed mice lacking the SOX17 enhancer associated with the risk of developing PAH in humans (8).

Using the CMap database, we identified compounds based on the prediction of their effects on gene expression, irrespective of whether these were traditional ‘on-target’ or ‘off-target’ effects when considering their known mechanisms of action. We prioritised BX-912 for *in vivo* testing due to its ability to restore the expression of SOX17 targets in PAECs and, particularly, RUNX1. In previous studies, we have shown that targeting RUNX1 attenuated PH in both Sugen-hypoxia and monocrotaline mouse models (39), as well as reversing PH in the more severe Sugen-hypoxia rat model (40), supporting its relevance to disease pathogenesis.

BX-912 was originally designed as a selective small molecule inhibitor of PDK1, and it antagonises ATP binding to PDK-1 (41). PDK-1 is a master kinase regulates the activity of multiple downstream kinases, including protein kinases A, C, N, and B (also known as AKT). AKT signalling is an established pathogenic pathway in PAH, with the AKT1 isoform required for the development of hypoxic pulmonary hypertension (42). PDK1 signalling also plays an important role in EC migration, with PDK1 loss hindering EC migratory potential and network formation and PDK1 overexpression leading to increased EC migration (43). PAEC dysfunction drives PAH pathology (44) and thus PDK1 inhibition by BX-912 might mitigate aspects of pathophysiological remodelling through improved EC function.

Prior studies have shown that mice with partial PDK1 knockout in endothelial cells (*PDK1^flox/+^: Tie2-Cre*) have an attenuated pulmonary hypertensive response to hypoxia (45). However, to our knowledge, no studies have demonstrated that PDK1 inhibitors can alleviate PH *in vivo*. It is possible that BX-912 also exerts effects on gene expression through PDK1-independent mechanisms, which remain to be determined, and therefore may be more efficacious than other PDK1 inhibitors not prioritised by our *in silico* analysis.

Our assumption from the offset was that we would identify drug compounds which affected expression of SOX17 target genes without directly affecting SOX17, as the CMap algorithm was blind to the fact that the genes we analysed were linked to SOX17. However, we found that BX-912 was able to increase *SOX17* expression in PAECs, though with minimal effects on protein levels under the conditions tested. This raises the possibility that at least some of its beneficial effect may be mediated through restoration of *SOX17* expression itself, thus facilitating the protective transcription factor activity of SOX17 in the pulmonary endothelium. Among the genes tested (including *SOX17*) *RUNX1* was the only target significantly affected by BX-912 treatment at 6 hours. This could indicate that BX-912 therapeutic activity is mediated through both SOX17-dependent and independent gene expression changes.

Beyond BX-912, we also identified four further new compounds, two of which reinstated the gene expression of the majority of SOX17 target genes we tested in PAEC. PIK-90, the compound which best corrected gene expression, is a phosphoinositide 3-kinases (PI3K) inhibitor, which acts by binding to the ATP site and inactivating PI3Kα/γ/δ, DNA-PK and PI3KC2s (46). The PI3K signalling pathway plays a crucial role in many biological processes such as angiogenesis, cell differentiation, proliferation and survival, and acts through various downstream signal mediators such as PKB/Akt, PDK1 and mTOR (47). PI3K signalling is also involved in neovascularization following ischemia, specifically via the effect of PI3Kγ isoform on EC proliferation and apoptosis (48). These processes are highly relevant to the development of PAH, which is characterised by hyper-proliferative and apoptosis resistant pulmonary ECs (44,49,50), supporting the prediction of PIK-90 as a potential therapeutic compound of PAH.

Most of the CMap-predicted compounds target pathways that are closely related or overlapping, including PDGF, mTOR, PI3K and PDK1 signalling. Studies have already demonstrated the added benefit of combined inhibition of mTOR and PDGF signalling in PH models, such as Sugen-hypoxia rats treated with sirolimus and imatinib (51), and have shown that PI3K signalling regulates crucial biological processes, including proliferation through downstream mediators such as PDK1 and mTOR (43,47). The precise involvement of SOX17 in these signalling pathways remains to be established, but these observations suggest potential benefit of combination therapeutic strategies targeting these pathways.

A logical question regarding the utility of the predicted compounds is whether they will be effective only in PAH patients with *SOX17* mutations. *SOX17* loss drives PAH development, and the distribution of both rare and common genetic variants indicates that the majority of PAH patients have some degree of genetic risk at this locus (∼three-fifths carry all four common risk alleles, and almost all carry at least one) (8).

Whether genetics will identify responders, or whether SOX17 represents a convergent pathway affected in all patients akin to BMPR2 signalling remains to be determined. However, we propose that *SOX17* is a promising therapeutic target, and that targeting this pathway may benefit the majority of patients rather than solely in those with rare pathogenic genetic variants. A parallel example is the recently approved PAH medication sotatercept, which was designed to correct the BMPR2 pathway dysfunction underlying 70-80% of heritable PAH. Notably, sotatercept was equally effective in patients with idiopathic PAH as in those with heritable PAH (52). This broad efficacy in PAH patients is likely due to the observation that BMPR2 signalling is disrupted and even in PAH patients without *BMPR2* variants, including increased pSMAD-2/3 signalling via the activin receptor IIA. We and others have shown that BMP9 enhances *SOX17* expression in PAECs (53), likely via BMPR2 signalling (54). More recent studies have shown that SOX17 expression is decreased in PAH patient and animal models of PH, and that TGFβ-1 suppresses SOX17 expression via increased pSMAD-2/3 activity (55). Thus, impaired SOX17 expression, whether or not patients are carriers of rare or common variants in *SOX17*, may contribute to PAH. Oral therapies that enhance SOX17 expression or modulate its downstream targets may therefore be a novel approach to treating most patients with PAH and could be particularly helpful in those with *SOX17* variants.

Whilst no doubt a useful resource, one of the biggest limitations of the CMap database is the exclusive use of cancer cell lines as opposed to primary cells for compound treatments. To address this limitation, in line with the directions of the CMap creators, we have used a detailed and robust signature for our analysis and subsequently measured the effect of the compounds on a number of up- and down-regulated genes in primary cells relevant to our disease of interest. Through our combination of multiple datasets to build the combined SOX17 transcriptomic signature, we observed an improvement in effectiveness (net benefit) of the compounds tested *in vitro* in this study compared to our previous one (16).

Besides being used to query the CMap, the SOX17 transcriptomic signature was also combined with ChIP-seq and PAH patient plasma proteomic data to validate the clinical relevance of the identified targets. Given the large surface area of the pulmonary endothelium, we expect that some of the proteins released by PAECs are detectable in circulating plasma, with a subset of these reflecting endothelial dysfunctions. Overlap of these datasets identified ten genes in which (i) mRNA levels are affected by SOX17 manipulation, (ii) plasma protein levels are affected by the genotype of *SOX17* signal 1 enhancer, and (iii) regulatory DNA are bound by SOX17. Amongst these candidate targets there are genes such as *EFNB2*, which plays a crucial role on EC network formation (56) and vessel sprouting (57), and *UNC5B*, which is implicated in vessel formation through the provision of repulsive cues (58) and the promotion of EC senescence (59). Loss of *EFNB2* and *UNC5B* affected viability and apoptosis in human PAECs, with *UNC5B* mimicking the effect of SOX17 loss, while *EFNB2* showed contradictory effects. This emphasises that the overall effect of SOX17 loss reflects the net result of multiple gene changes, with some alterations being protective as others are harmful. This balance in PAH is clearly detrimental, and targeting the appropriate downstream pathways will be important for fine-tuning strategies aimed at restoring homeostatic SOX17 function.

The function of SOX family members is highly affected by the binding of co-factors, yet there is very little information on the co-factors of SOX17 in vascular endothelium. For this reason, we utilised our ChIP-seq dataset to systematically search for SOX17 co-factors, which indicated enriched binding of the TF ERG and other ETS members in proximity to SOX17 in PAECs. ERG is a crucial endothelial TF (60,61), and it has been shown that pathologic high shear stress (HSS) negatively regulates ERG, leading to endothelial to mesenchymal transition (EndoMT) (60). Recent studies have shown that *SOX17* overexpression can inhibit EndoMT (55). ERG levels are reduced in the lungs of PAH patients and increased monolayer permeability follows ERG loss (61), mimicking *SOX17* knockdown (16). These findings imply that SOX17 could potentially cooperate with ERG in the regulation of PAEC function in maintaining pulmonary vasculature homeostasis.

## Conclusions

By identifying gene targets of the PAH risk gene *SOX17*, we predicted a novel therapeutic agent, BX-912, which reverses PAEC gene expression and functional changes associated with SOX17 loss and prevents PH development in a mouse model recapitulating common genetic risk of PAH. These findings prioritise BX-912 and its targets for further development and provide proof-of-concept that targeting omics-derived disease signatures can offer a promising therapeutic strategy in PAH.

## Funding

CJR is supported by British Heart Foundation (BHF) Basic Science Research fellowship (FS/SBSRF/22/31025) and Academy of Medical Sciences Springboard fellowship (SBF004\1095). This work was also supported by BHF Project grant PG/19/17/34275 and BHF Imperial CRE PhD Studentship to EV FS/19/57/34894A and BHF CRE award RE/18/4/34215. This work was supported by the National Institute for Health and Care Research (NIHR) BioResource which supports the UK National Cohort of Idiopathic and Heritable PAH; BHF SP/12/12/29836 and the UK Medical Research Council (MR/K020919/1). NIHR BRC awards to Imperial College and Cambridge also supported this study. This work was further supported by NIH/NHLBI R01 HL158841 (Olin Liang & James Klinger), NIH/NHLBI R01 HL174007 (Corey Ventetuolo & Olin Liang), NIH/NHLBI T32HL160517 (Olin Liang & Bedia Akosman), NIH/NIGMS P30 GM145500 (Olin Liang), NIH/NIGMS P20 GM103652 (James Klinger & Olin Liang), Brown Physicians Incorporated Academic Assessment Research Awards (Olin Liang & James Klinger). IC is recipient of a Sir Henry Dale Fellowship jointly funded by the Wellcome Trust and the Royal Society (224662/Z/21/Z).

## Disclosures

CJR, MRW, LZ, EV, RW, OL and JK have filed a patent for the use of BX-912 as a therapeutic for PAH. LH discloses Honoraria for lectures (Merck Sharp & Dohme Ltd, Janssen, AOP, Ferrer), Honoraria for scientific advisory boards / steering committees (Janssen, Gossamer Bio, Merck Sharp & Dohme Ltd, United Therapeutics, Apollo Therapeutics, Liquidia, Morphic, Keros Therapeutics, Biologix, Pulmovant), Research support (Merck Sharp & Dohme Ltd.). CEV discloses personal fees from Merck & Co., Janssen Pharmaceuticals, and Pulmovant outside of the submitted work and Patent application BM-2024-052; 15024-398PC0 pending to None. Her institution has received fees for the conduct of clinical trials from Merck & Co., Pulmovant, Tenax, Gossamer and United Therapeutics.

## Supplemental Material

### Supplementary files

**Supp Excel 1 - SOX17 RNAseqs DEG.** List of DEGs derived from the RNA-seq analyses following SOX17 knock-down and overexpression (p<0.05 & Log_2_FC>|0.25|).

**Supp Excel 2 – GREAT gene annotation.** Genes bound by SOX17 in ChIP-seq annotated by GREAT and expressed in PAECs

**Supp Excel 3 - Primary motifs.** Primary known and *de novo* motif enrichment analysis on SOX17 ChIP-seq peaks in PAECs

**Supp Excel 4 - Primary & coenriched motifs.** Primary and coenriched known and *de novo* motif enrichment analysis on SOX17 ChIP-seq peaks in PAECs.

### Supplementary Figures

**Supplementary Figure 1.** Levels of SOX17 overexpression in PAECs at 48 hours.

**Supplementary Figure 2.** Differentially expressed genes following SOX17 overexpression with Adenoviral vectors (MOI 1).

**Supplementary Figure 3.** Over-representation analysis for enriched functions and pathways following SOX17 overexpression (o/e) (Adv MOI 5).

**Supplementary Figure 4.** Over-representation analysis for enriched functions and pathways following SOX17 overexpression (o/e) (Adv MOI 1).

**Supplementary Figure 5.** Effect of SOX17 overexpression on novel downstream targets

**Supplementary Figure 6.** Binding of SOX17 at putative targets’ regulatory regions.

**Supplementary Figure 7.** Downregulation of *EFNB2* and *UNC5B* in PAECs.

**Supplementary Figure 8.** Smooth muscle actin staining in WT or SOX17 enhancer knockout mice exposed to mild Sugen-Hypoxia plus BX-912.

